# Neutralizing Activity of Cervicovaginal Secretions against Herpes Simplex Virus is Mediated by Mucosal IgG and Viral Glycoprotein E and Adversely Impacted by Vaginal Dysbiosis

**DOI:** 10.1101/2025.05.15.654401

**Authors:** Aakash Mahant Mahant, Valerie Fong, Matthew Gromisch, Richard Hunte, Ian Michael, Jennifer T. Aguilan, Kerry Murphy, Marla J. Keller, Betsy C. Herold

**Author notes:** Address correspondence to Betsy C. Herold,. Aakash Mahant and Valerie Fong contributed equally to this work. Author order was determined on the basis of seniority.

## Abstract

Genital herpes simplex virus (HSV) recurrences are more common in women with bacterial vaginosis (BV). Prior studies demonstrated that genital tract secretions exhibit variable neutralizing activity against HSV, independent of serostatus, but the relationship of this activity to the vaginal microbiome and underlying mechanisms have not been defined. To test the hypothesis that cervicovaginal antiviral activity is lower in women with BV, we took advantage of cervicovaginal lavage (CVL) available from two studies conducted among women with symptomatic BV and healthy controls. CVL obtained from women with BV had significantly less antiviral activity than controls (p< 0.001). Inhibitory activity correlated negatively and most strongly with Shannon diversity index (p<0.0001). The innate activity did not differ comparing HSV-seropositive versus seronegative participants and no HSV-specific antibodies were detected in CVL. Activity was enriched in the immunoglobulin fraction but was lost when IgG (but not IgA) was depleted. Increasing doses of an anti-glycoprotein E (gE) monoclonal antibody overcame the neutralizing activity, suggesting that interactions between the Fc region of IgG and gE, a viral Fc gamma receptor (FcγR), contribute. Consistent with this notion, CVL had less HSV inhibitory activity against a gE-null virus. Glycan analysis demonstrated a decrease in mature glycans in IgG from CVL with low antiviral activity and treatment of CVL with peptide N-glycanase F, which cleaves N-glycans in IgG, resulted in a loss of HSV inhibitory activity. We speculate that glycosidases elaborated by anaerobic bacteria cleave Fc glycans, resulting in decreased affinity for gE and a reduction in protective activity.

IMPORTANCE: This study provides a mechanistic link for the increased risk of HSV infection and replication in the setting of symptomatic bacterial vaginosis and asymptomatic vaginal dysbiosis. Independent of Fab antigen specificity, the Fc region of mucosal IgG may neutralize HSV by binding to glycoprotein E, a viral Fc receptor. Vaginal dysbiosis leads to a loss of Fc glycans and a concomitant decrease in this innate antiviral activity. These findings suggest that viral Fc receptors, previously thought to function only in immune evasion, may also play a protective role. The results highlight the importance of developing and implementing strategies to protect against vaginal dysbiosis.

## Introduction

Herpes simplex virus type 1 and type 2 (HSV-1 and HSV-2) infections are major global health burdens. Both types cause genital mucocutaneous disease with the potential risk of perinatal transmission and may be linked to HIV acquisition and latency reversal(1–3). Specifically, the high prevalence of HSV-2 in sub-Saharan Africa (∼70%) has been implicated as a major cofactor in the HIV epidemic (4–6). The impact of infection is life-long because the viruses establish latency with variable frequency of subclinical or clinical reactivations (7). Genital HSV infections are more common in young women compared to men, which contributes to the risk of perinatal transmission. The reasons for the increased risk of genital HSV in young women are complex and likely reflect biological and behavioral factors. Among the associated biological factors is bacterial vaginosis (BV). Epidemiological studies demonstrate that women with BV have a greater incidence of genital herpes (8), although the mechanisms are not clear.

In prior studies, we found that genital tract secretions collected either by cervicovaginal lavage (CVL) or vaginal swab from HSV seropositive or seronegative women had variable anti-HSV activity and inhibited viral entry and plaque formation when preincubated with virus in vitro(9–11). This “neutralizing” activity correlated with the concentrations of several mucosal immune molecules including Immunoglobulin (Ig) A, IgG, human neutrophil peptides (HNP1-3), secretory leukocyte protease inhibitor (SLPI) and select proinflammatory cytokines and chemokines (10, 11). The potential for this innate activity to contribute to viral protection was suggested by the finding of reduced disease in mice infected intravaginally with HSV-2 that had been preincubated with human CVL possessing HSV neutralizing activity(9). However, precisely what mediates this activity and whether it is modulated by the vaginal microbiome has not been studied.

BV impacts the concentration of several molecules that are correlated with the anti-HSV activity. For example, bacteria that dominate in the setting of BV have increased glycosidase activity, which may result in degradation of glycans expressed by Igs and alter Ig function (12, 13). Based on these findings, coupled with the epidemiological link between BV and HSV, we hypothesized that the anti-HSV activity of CVL may be reduced in women with BV or vaginal dysbiosis. To test this hypothesis, we took advantage of CVL previously obtained from a study of US women with symptomatic BV before and after metronidazole treatment (14) and, from a second study that enrolled US women with symptomatic BV and controls without BV (manuscript in preparation). We found that the CVL neutralizing activity was reduced in women with clinical BV and, specifically among those with a vaginal microbial community dominated by anaerobes (15). We conducted CVL enrichment and depletion studies, which demonstrated that the anti-HSV activity mapped to the IgG-enriched fraction but was independent of HSV serostatus. The neutralizing activity was overcome by addition of a monoclonal antibody (mAb) targeting HSV glycoprotein E (gE), which is a viral Fcgamma receptor (FcγR), and showed decreased activity against a gE null virus, suggesting a previously unrecognized protective role for IgG Fc and HSV gE interactions. We speculate that the cleavage of Fc glycans on IgG by glycosidases elaborated by vaginal anaerobes results in decreased affinity for gE and contributes to decreased HSV neutralizing activity of the CVL observed in the setting of BV. Consistent with this notion, we found differences in glycans in samples obtained from participants with high versus low anti-viral activity and a decrease in neutralization when CVL samples were subjected to enzymatic digestion of N-glycans.

## RESULTS

### BV is associated with significantly less anti-HSV activity in genital tract secretions

CVLs were evaluated for HSV antiviral activity using stored samples from two prior studies. In the first study, CVLs were obtained from 20 US women at the time of BV diagnosis (Day 0) and approximately one week (Day 14) and one month (Day 35) following a 7-day course of twice daily oral metronidazole (14). Demographic data is shown in **Table 1**. CVLs obtained at the time of BV diagnosis reduced HSV plaque formation by a median [IQR] of only 5.8% [-8.9% - 47.1%]. There was a small but non-significant statistical increase in the anti-HSV activity of CVL obtained one week or four weeks following metronidazole treatment 14.1% [-2.6% - 31.3%] and 15.2% [-3.7% - 41.8%], respectively (**Figure 1A**). This study did not include a control (non-BV) population, but the results differed from prior studies that only enrolled women without BV. For example, CVL inhibited HSV plaque formation by a mean of 57% in cycling women and by a mean of 36% in women on hormonal contraception in a prior study that collected 3-8 weekly samples (11). The anti-HSV activity was stable throughout the study period, but, notably, the microbiome was not assessed.

**Table 1.** Demographics and vaginal microbial community state types (CST) for the study population

|  | Study A<br>(BV cases)<br>(n=20) | Study B<br>(BV cases)<br>(n=18) | Study B<br>(Controls)<br>(n=15) |
| --- | --- | --- | --- |
| Age (median [IQR] years) | 39 [30, 42.75] | 25.5 [20.5, 29.5] | 24 [21, 25] |
| Race (White: Black: Other) | 5: 13: 2 | 2: 6: 10 | 7: 4: 2 |
| Ethnicity (Hispanic: Non-Hispanic) | 9: 11 | 8: 10 | 6: 7 |
| Hormonal contraception (Y: N) | 4: 16 | 9: 9 | 8: 7 |
| Community state types (I: II: III: IV: V) | 3: 1: 1: 14: 0 | 3: 0: 0: 15: 0 | 9: 0: 3: 3: 0 |

CSTI, *L. crispatus*; CSTII, *L. gasseri*; CSTIII, *L. iners*; CSTIV, anaerobic dominance; CSTV, *L. jensenii*. One Study A participant had missing microbiome data.

**Figure 1:**
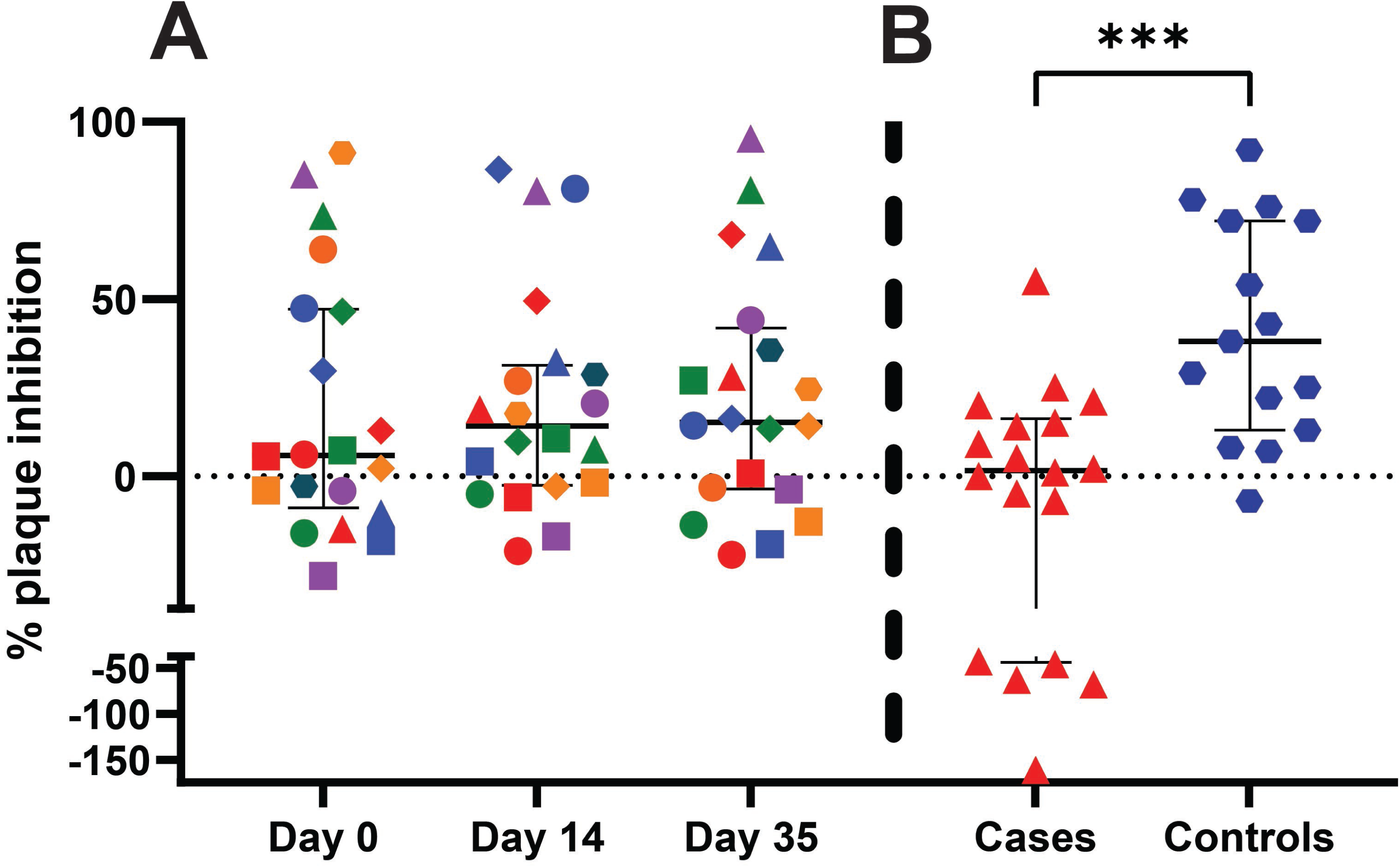
BV is associated with less anti-HSV activity in genital tract secretions. (A) HSV activity was measured by plaque reduction assay in CVL of US women who were diagnosed with symptomatic BV at Day 0 and treated with metronidazole and followed at Day 14 and 35. Each participant is color-coded, and results are shown as median (IQR) percent inhibition relative to virus treated with control buffer. (B) CVL activity was measured in CVL obtained from participants in a second study (Day 0 only) that recruited US women with symptomatic BV (cases) and controls and compared by unpaired t-test (***p<0.001).

To address the differences observed between these two studies, we took advantage of CVL available at enrollment from a second study that included 18 participants with BV diagnosed by demonstrating at least 3 of 4 Amsel criteria and 15 age-matched controls without BV (Table 1 and manuscript in preparation). Consistent with the first study, CVL from participants with BV inhibited HSV by only 1.5% [-43.8% - 16.3%] with a few CVL increasing HSV infection. In contrast, CVL from the controls (n=15) inhibited plaque formation by 38.0% [13.0% - 72.0%], p< 0.01) (**Figure 1B**). The number of women on hormonal contraception was similar in the cases and controls and there were no differences in the anti-HSV activity comparing those who were or were not on hormonal contraception.

### Anti-HSV activity of CVL correlates negatively with biomarkers of vaginal dysbiosis

To explore the relationship between BV and anti-HSV activity, we correlated the percent inhibition from participants in both studies (only Day 0 CVL from the first study and CVL from cases and controls from the second study) with the Nugent score, Shannon diversity index (SDI), and community state types (CST) (15); the latter were determined using 16srRNA sequencing data with CST defined by having 50% or greater of the dominant species (14). The HSV inhibitory activity (n=51; 2 of the controls had incomplete data and were excluded) correlated negatively with all three measures of the microbiome (Nugent, SCC= -0.48, p< 0.0001; SDI, SCC= -0.43, p=0.002; and CST, SCC=-0.46, p< 0.001) (**Figure 2A**). Similar to prior studies, the inhibitory activity also correlated modestly and positively with IL-6 (SCC 0.31, p=0.026), IL-8 (SCC 0.38, 0.014), CXCL9 (SCC 0.45, p=0.001), CXCL10 (SCC= 0.38, p= 0.006), MIP-1α (SCC=0.56, p=0.001) and IgA (r=0.45, p=0.001), but only trended to correlate with IgG (r=0.25, p=0.08). There were 15 participants with CSTI (*L. crispatus* dominant) and 32 with CSTIV (anaerobe dominant) but only one with CSTII (*L. gasseri*), two with CSTIII (*L. iners*) and none with CSTV (*L. jensenii*). There was significantly less anti-HSV activity in CVL obtained from women with CSTIV vs CSTI and the five CVL that increased HSV infection (plaque forming units, pfu) had CSTIV microbiomes (**Figure 2B**, p<0.001).

**Figure 2:**
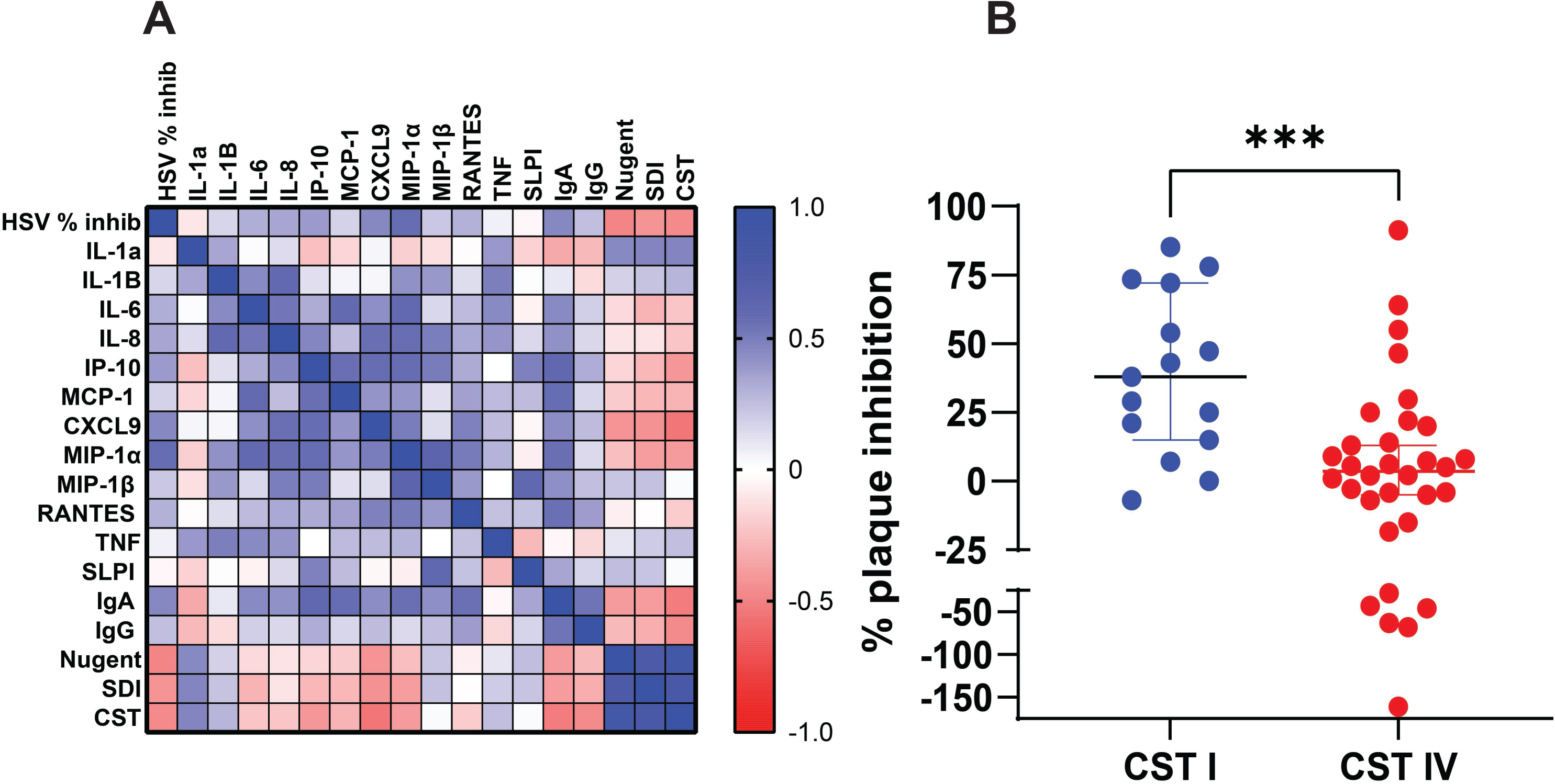
Anti-HSV activity correlates negatively with biomarkers of vaginal dysbiosis. (A) Spearman rho correlation matrix of HSV inhibitory activity, biomarkers of vaginal microbiome and concentrations of mucosal immune mediators using data from n=51 participants (Visit 1 only) from both studies. (B) HSV percent inhibition comparing samples from participants with CST I and CST IV vaginal microbiomes. Results are shown as median (IQR) and compared by Mann-Whitney test (***p<0.001).

### Anti-HSV activity maps to immunoglobulins

To identify molecules within CVL that might contribute to the anti-HSV activity, we analyzed eight samples with variable inhibitory activity (**Figure 3**). Each CVL sample was applied to a Protein L column, which binds Igs and the flow through and eluant were collected, protein concentration quantified, and material tested for anti-HSV activity by plaque assay after diluting to control for the protein concentration (0.5 mg/ml). The anti-HSV activity was significantly greater in the Protein L eluant compared to the flow through (p< 0.01, Wilcoxon test), suggesting that it maps to Igs.

**Figure 3:**
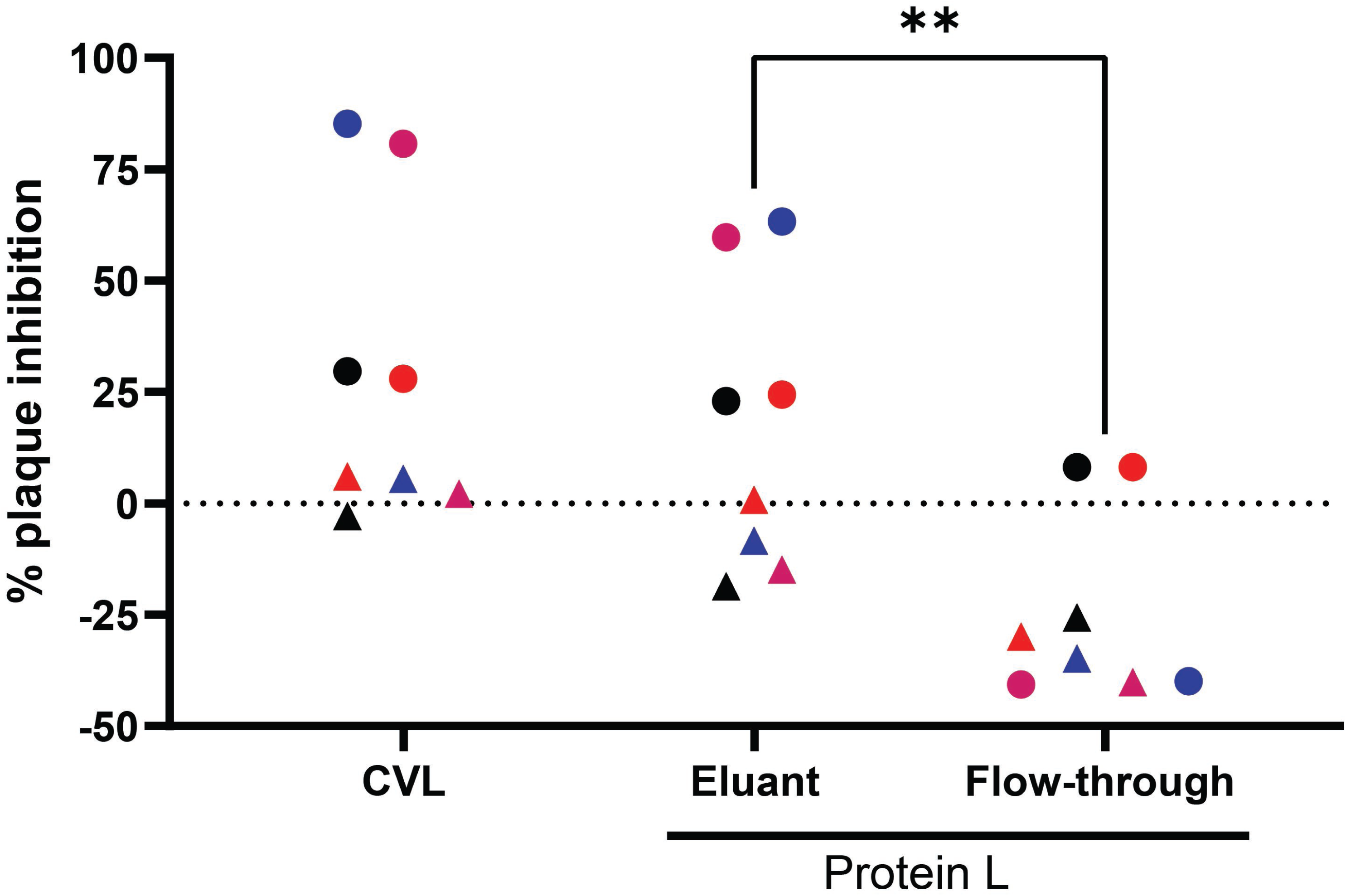
Anti-HSV activity maps to immunoglobulins. CVL (n=8) were applied to Protein L columns to isolate immunoglobulins and non-immunoglobulin proteins. The samples were diluted to a final protein concentration of 0.5 mg/ml and the anti-HSV activity was measured in starting material, flow through and eluant by plaque assay. Results are presented as the mean (duplicate wells) percent inhibition of plaque formation relative to virus treated with control buffer. Each CVL is color-coded; circles were samples with anti-HSV activity and triangles are samples with little or no anti-activity. The percent inhibition is compared between eluant and flow through by paired t-test (**p<0.01).

To confirm this observation and to assess whether it was associated with IgG or IgA, we took advantage of CVL from 5 additional participants with high activity (73.7 ± 4.9%) and 4 with low activity (31.2 ± 6.8%) (**Table 2**); these nine samples were selected based on number of CVL aliquots available for mechanistic studies. We depleted the CVL’s of IgA, IgG, or both using antibody magnetic beds. The protein concentration was quantified, and samples diluted to control for protein concentration (0.5 mg/ml) before testing in the HSV plaque assay. For the 5 CVLs with high anti-HSV activity, the activity was retained in the IgA depleted material (71.3 ± 4.7%) but significantly reduced when IgG (17.4 ± 3.9%) or both IgG and IgA (8.8 ± 1.9%) were depleted (p<0.0001, ANOVA with Dunnett’s multiple comparisons) (**Figure 4A**). Depletion of IgA and IgG was confirmed by ELISA (**Figure 4B and 4C**). To control for specificity of the depletion, we also measured the concentration of HNP1-3, which was unaffected by the depletion (**Figure 4D**). Notably, there was no difference in the concentration of IgA or IgG in CVL with high vs low activity, suggesting that quantity does not explain the functional differences. Possibly the small amount of activity retained in the IgG depleted fractions reflects contributions from other molecules including HNP1-3, which in vitro, has anti-HSV activity (9).

**Table 2.** Demographic data, HSV serostatus and microbiome community state types (CST) for nine control participants with CVL aliquots available for depletion and glycan studies.

| ID | Age | Race/<br>Ethnicity | HSV<br>serostatus | HSV %<br>inhibition | Microbiome Community State Types |
| --- | --- | --- | --- | --- | --- |
| CVL with high HSV activity |  |  |  |  |  |
| 1 | 21 | Bl/His | HSV2+ | 76% | CSTIII (*L. iners*) |
| 2 | 19 | Asian | HSV1+ | 72% | CSTIII (*L. iners*) |
| 3 | 25 | White | Seronegative | 81% | CSTIII (*L. iners*) |
| 6 | 24 | White | Seronegative | 70% | CSTI (*L. crispatus*) |
| 9 | 24 | White | Seronegative | 69% | CSTI (*L. crispatus*) |
| CVL with low HSV activity |  |  |  |  |  |
| 7 | 23 | White | HSV1+ | 40% | CSTIV (Anaerobe dominance) |
| 8 | 25 | Bl/His | Seronegative | 27% | CSTI (*L. crispatus*) |
| 4 | 21 | Asian | HSV1+/HSV2+ | 25% | CSTIV (Anaerobe dominance) |
| 5 | 21 | Bl/His | Seronegative | 32% | CSTIV (Anaerobe dominance) |

**Figure 4:**
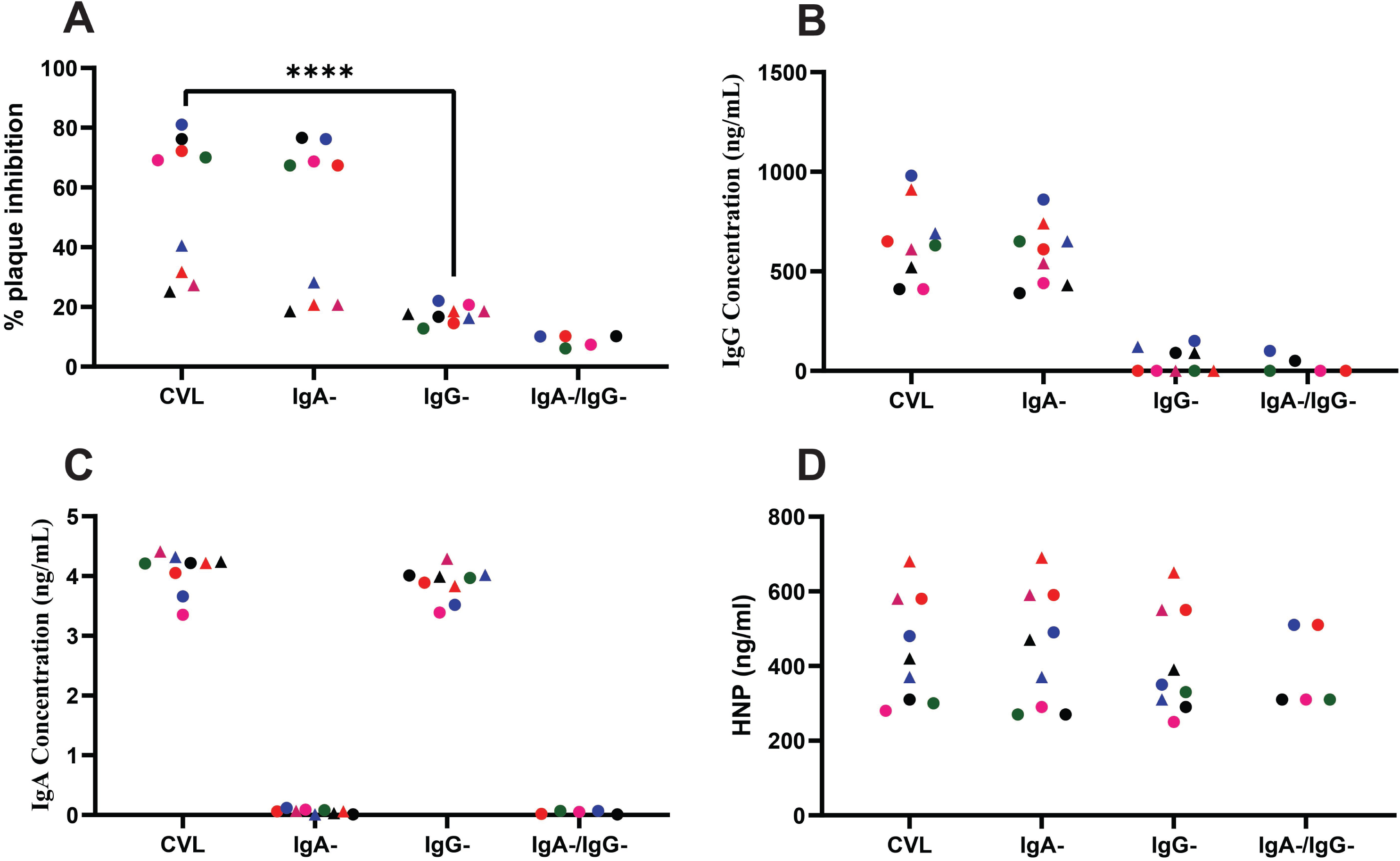
Anti-HSV activity was enriched in CVL samples depleted with IgA but lost when IgG was depleted. (A) Percent inhibition of HSV plaque formation, (B) IgG concentrations, (C), IgA concentrations, and (D) HNP1-3 concentrations were measured in CVL or CVL depleted of IgA, IgG, or both from n=9 different participants; five with high (circles) and four with low (triangles) anti-HSV activity. Each participant’s CVL is color-coded. The percent inhibition of HSV plaque formation was compared by ANOVA with Dunnett’s multiple comparisons (***p<0.0001).

To determine if the anti-HSV activity mediated by the IgG enriched CVL samples was mediated by HSV-specific antibodies, we compared antiviral activity in HSV seropositive vs seronegative participants and measured HSV-binding antibodies in the CVL by ELISA. Serum was available from 18 BV cases and 9 controls to test for HSV serostatus (HerpeSelect immunoblot). All but two of the BV cases were HSV seropositive compared to 4/9 controls (Chi square, p=0.09) and there was no difference in CVL anti-HSV activity comparing seronegative vs seropositive participants (**Figure 5A**). Moreover, using CVL from the same nine participants whose samples were used for the IgG/IgA depletion studies, we detected little or no HSV specific IgG in the CVL by ELISA (**Figure 5B**). Serum from the same participants was included in the ELISA assay as controls and matched the HerpeSelect immunoblot results (**Figure 5C**). There was also no HSV-specific IgA detected in the CVL by ELISA (**Figure S1**).

**Figure 5:**
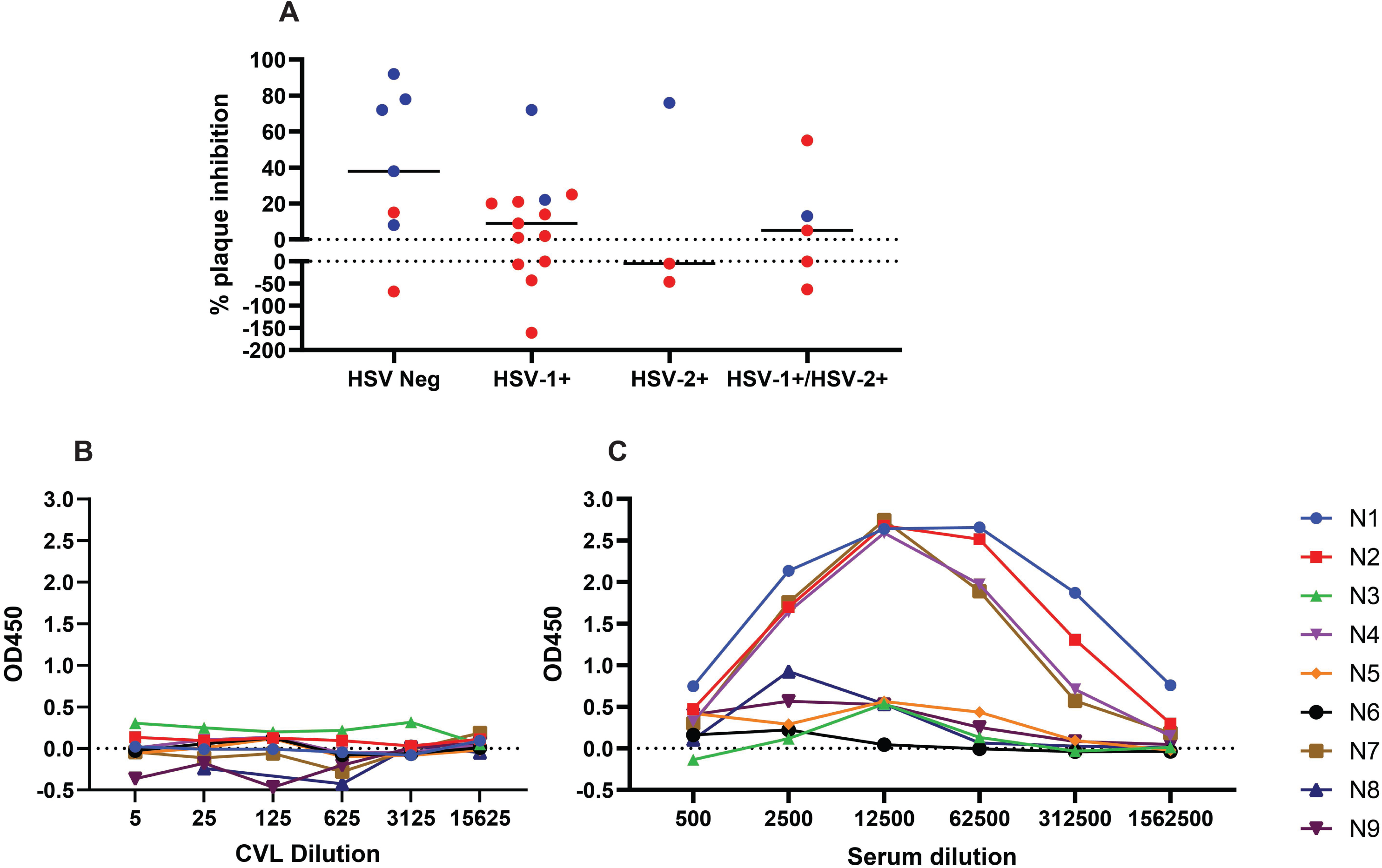
CVL anti-HSV activity is independent of HSV-specific antibodies. (A) Serum was available from n=19 BV cases (red circles) and n=9 controls (blue circles) (Study B) to assess HSV serostatus (FOCIS immunoblot assay). The percent inhibition of HSV plaque formation was compared for participants who were HSV seronegative, HSV-1 seropositive, HSV-2 seropositive or dually HSV-1 and HSV-2 seropositive. (B) HSV IgG was measured in serial dilutions of CVL or (C) serum by ELISA using samples from indicated controls (Table 2).

### Interactions between IgG and the viral Fc receptor, glycoprotein E, contribute to the antiviral activity of CVL

Because the anti-HSV activity of CVL IgG was independent of HSV specific antibodies and did not correlate with IgG concentration, we speculated that some other feature(s) of the IgG contributed to its ability to inhibit HSV infection. We hypothesized that the Fc component of CVL IgG, independent of its antigenic target, binds to HSV gE, a known viral FcγR. To test this notion, we added increasing concentration of a non-neutralizing gE mAb or, for comparison, a non-neutralizing gB mAb, to five different CVLs with HSV neutralizing activity prior to conducting the plaque assay. The anti-gE but not the anti-gB mAb overcame the CVL inhibitory activity in a dose dependent manner (**Figure 6A**). Similar results were obtained whether we used the intact mAb or the Fab fragment (**Figure S2**). Consistent with these findings, we found significantly less CVL anti-HSV activity against a gE-deleted compared to wild-type virus (p<0.001, paired t-test, n=5) (**Figure 6B**).

**Figure 6:**
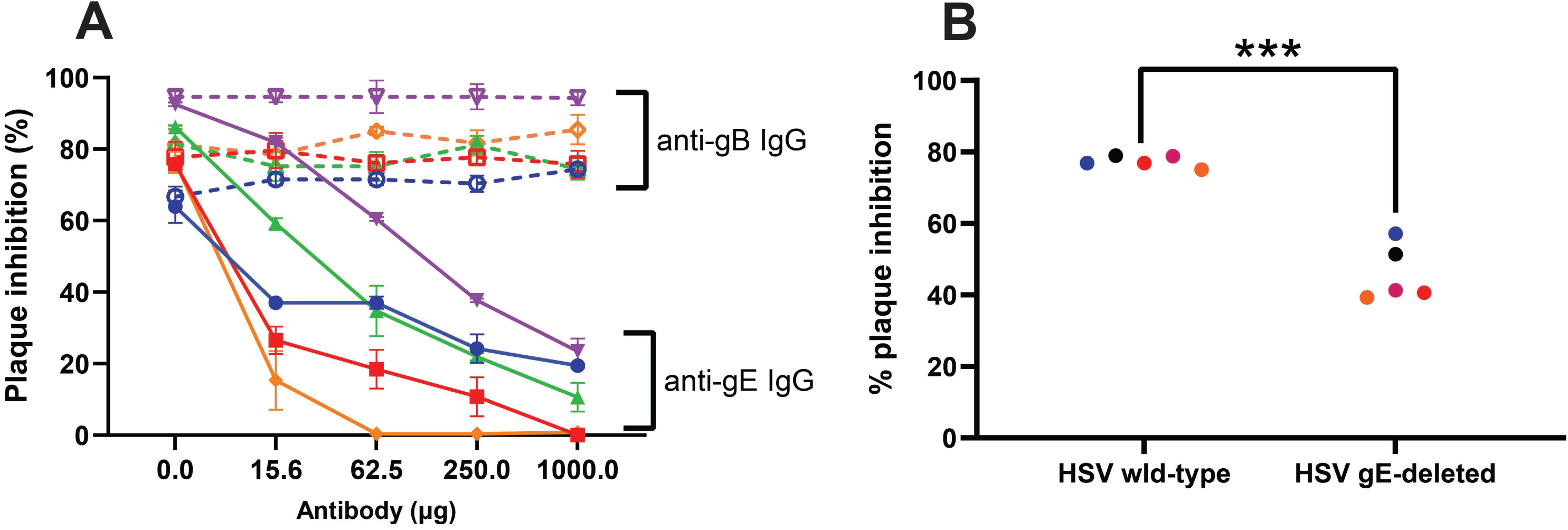
Interactions between IgG and the viral Fc receptor, glycoprotein E, contribute to the antiviral activity of CVL. (A) Plaque assays were performed with CVL from the n=5 controls with high anti-HSV activity in the presence of increasing concentrations of an anti-gE (closed symbols and line) or anti-gB monoclonal antibody (open symbols and line). Results are shown as percent inhibition relative to infection in the presence of control buffer. (B). Anti-HSV activity was measured by plaque assay with HSV-1 wild-type or an HSV-1 gE-deleted virus and the percent inhibition was compared between the two viruses (paired t-test, **p<0.001). CVL from participants is color coded as in **Figure 5**.

### CVL IgG with low anti-HSV activity express fewer mature glycans

Changes in the glycosylation pattern of the Fc domain of IgG modulate its affinity for cellular FcγRs (16). We hypothesized that the association between the vaginal microbiome and the antiviral activity of CVL may be attributed, in part, to enzymes elaborated by anaerobic bacteria that degrade N-glycans. To test this hypothesis, we isolated and characterized the N-glycans in IgG enriched CVL (n=9, **Figure 4**). The IgG glycans in the CVL with high HSV inhibitory activity expressed more mature glycans (m/z >1836) compared to IgG glycans evaluated from participants with low inhibitory activity (38.17% ± 34% vs 10.4% ± 7.5%, p<0.01, respectively, **Figure 7** and S3). However, these differences did not segregate completely by CST as the three samples with the most mature glycans came from women with CSTIII (*L. iners* dominant) microbiomes. A more detailed analysis of the glycans showed that samples with CSTIII had higher amounts of bisected N-acetyl glucosamine (GlcNAc) with mono- or di-galactose glycans; bisected GlcNAc Fc glycans are associated with increased binding affinity for FcγRs (17) (**Figure S3**). Specifically, all five of the samples with anti-HSV activity had G1F0B (m/z 2111.2) glycan detected whereas this glycan was not detected in the noninhibitory samples (**Figure S3**).

**Figure 7:**
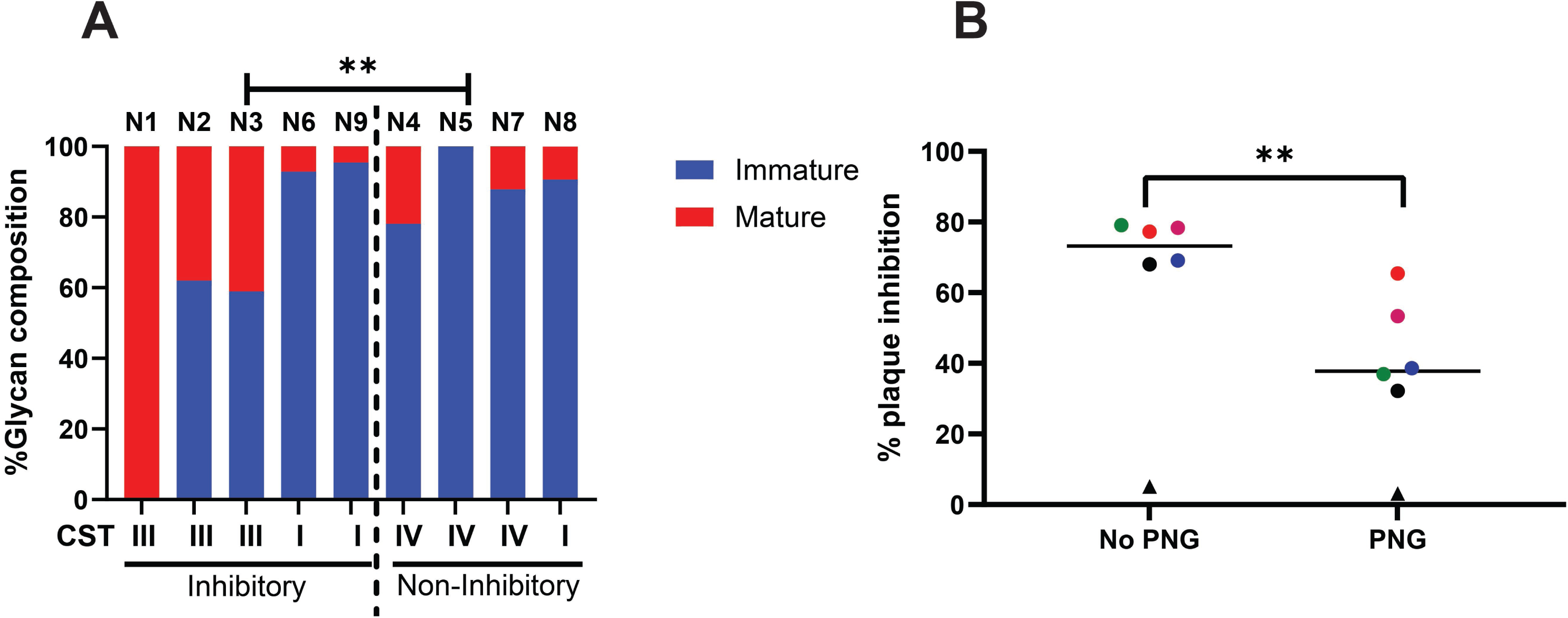
Differences in glycans in samples with high versus low HSV inhibitory activity. (A) N-linked glycans were isolated characterized by MALDI-TOF mass spectrometry from the 5 CVL with high vs 4 with low HSV inhibitory activity and dichotomized into mature (m/z >1836) vs immature and compared using 2-way ANOVA (**p<0.01). The community state types (CST) are indicated below each bar. (B). CVLs from n=6 participants (5 with high anti-HSV activity and 1 with low anti-HSV activity) were subjected to enzymatic N-glycan digestion with PNGase-F and the anti-viral activity assessed by plaque reduction assay. Results are presented as percent inhibition (relative to control buffer) and compared by paired t-test for the 5 samples with anti-viral activity (**p<0.01).

To further explore whether N-glycans modulate the anti-HSV activity in CVL, we subjected CVLs (n=6, 5 with inhibitory activity and one without) to enzymatic digestion with PNGase F, which cleaves N-linked oligosaccharides. Treatment with PNGase F resulted in a significant reduction in the anti-HSV activity of the inhibitory CVLs (74.37±5.4 vs 45.32±13.8, p<0.01), (**Figure 7B**).

## Discussion

Results of this study support the hypothesis that vaginal dysbiosis is associated with less “innate” anti-HSV activity in cervicovaginal secretions. Using clinical samples from two different studies, we found that the neutralizing activity of CVL correlated most strongly and negatively with biomarkers of vaginal dysbiosis even in the absence of clinical BV. These findings may provide a biological basis for the clinical observations that BV is associated with increased HSV acquisition and shedding(18, 19). For example, in a study of women with acute HIV infection (CAPRISA 002), BV (diagnosed by Nugent score) was associated with an increased risk of HSV-2 shedding in vaginal swabs (adjusted increased hazard ratio 2.17 [95% CI 1.15 - 4.10]) (20). Previously proposed mechanisms that could contribute to these epidemiological findings include a compromised epithelial barrier and mucosal inflammation, which may facilitate HSV acquisition and replication. Our studies provide an additional new mechanism mediated by mucosal IgG binding to viral gE.

While the ability of genital tract secretions to inhibit HSV infection did not correlate significantly with the concentration of IgG, enrichment and depletion studies mapped the antiviral activity to the IgG fraction. HSV specific antibodies were not detected in the CVL and HSV serostatus was not associated with the antiviral activity, suggesting that it is neither the quantity nor the antigenic specificity of the IgG that determines its inhibitory action. These observations led us to hypothesize that the activity might be mediated by interactions between the Fc region of IgG and the HSV FcγR mimetic, gE. Interactions between gE (or more potently, the heterodimer of gE-gI) and the Fc region of HSV-specific IgG have been studied primarily as an immune evasion strategy. HSV-specific antibodies bind to viral glycoproteins (e.g., gB, gC or gD) via their Fab regions but also bind to gE (or the gE-gI complex) via their Fc. This Fc-gE interaction functions as an immune evasion strategy because it interferes with Fc-mediated activities such as antibody-dependent cellular cytotoxicity (ADCC) (21, 22). Our results suggest a different paradigm as illustrated in **Figure 8**. We propose that, when HSV-specific antibodies are absent or present at low levels, interactions between the Fc of non-HSV specific mucosal IgG and viral gE neutralize the virus to protect the host. This Fc-gE protective mechanism is supported by the finding that addition of an anti-gE mAb (but not a gB mAb) overcame the CVL mediated inhibitory activity and the finding that CVL was less effective at inhibiting infection by a gE null virus. This may explain why the “innate” anti-HSV activity of the CVL was enriched in the IgG rather than IgA fraction as viral (and cellular) FcγRs bind IgG whereas IgA primarily binds FcαR1.

**Figure 8.**
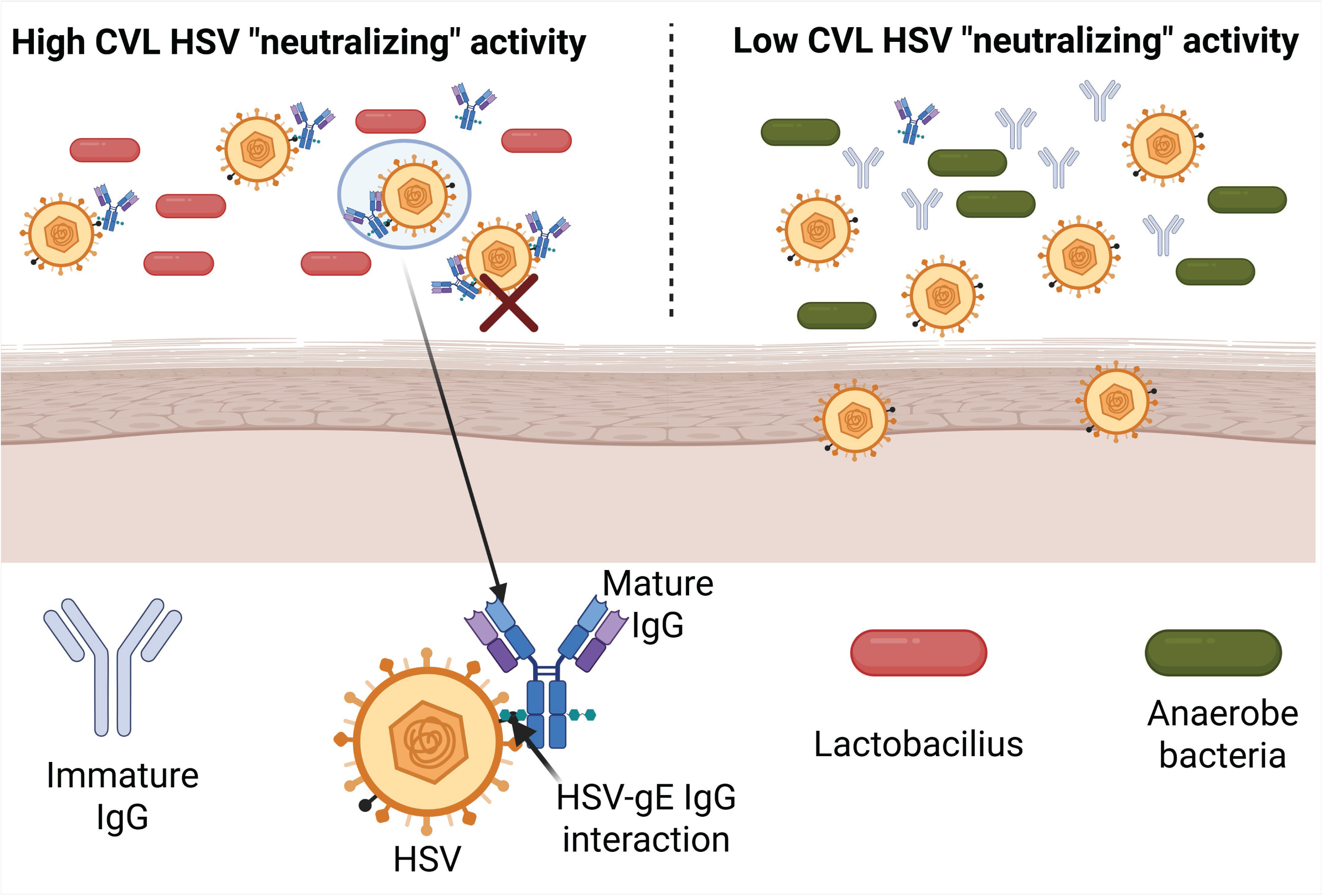
Model of how non-HSV specific IgG interacts with gE to reduce viral infection. IgG present in genital tract secretions binds via its Fc to HSV glycoprotein E on viral particles and sterically inhibits viral infection. The affinity of the Fc for gE is influenced by N-glycans. However, when the Fc glycans are cleaved by enzymes secreted by vaginal anaerobic bacteria, the affinity for gE is decreased and there is a loss of this neutralizing activity resulting in productive viral infection.

Whether the immune evasive or immune protective effects of gE-Fc interactions dominate may depend on the balance between HSV-specific and nonspecific IgG present at the site of viral infection. HSV-specific antibodies are detected at much lower levels in genital tract secretions compared to the serum in seropositive individuals in the absence of viral shedding, as evidenced here by the inability to detect HSV-binding IgG by ELISA in CVL in HSV seropositive participants. Thus, the Fc-gE interaction may provide an immediate protective role prior to the transport of HSV-specific antibodies and cytolytic T cells to the sites of viral replication. The role of gE may shift to immune evasion with the increase in mucosal HSV-specific antibodies but is ultimately overcome by the combination of protective antibodies and cytolytic T cells that lead to viral clearance.

The notion that HSV-specific antibodies are recruited into sites of viral replication is supported by studies in both mice and humans. For example, only low levels of HSV-specific antibodies were detected in the skin of mice that had been vaccinated with a single-cycle vaccine that induces high titer serum antibodies, but the antibody levels rapidly increased as early as two days after viral challenge in vaccinated but not control mice. The antibodies recovered from the skin were predominantly IgG (23). Similarly, B cells and antibody-secreting cells were detected in skin biopsies obtained from patients during symptomatic HSV-2 recurrences and colocalized with CD4+ T cells. There was also an increase in the concentration of HSV-2 specific antibodies in the affected skin but not the serum. In contrast, HSV-specific antibodies and antibody-secreting B cells were rarely detected in biopsies of unaffected skin in the seropositive patients (24).

This innate neutralizing activity was significantly less in CVL obtained from women with clinical BV and, specifically, among women with an anaerobic dominant vaginal microbiome. This low level of activity was observed in two independent studies of women with symptomatic BV, although only one study included controls without BV. Precisely why vaginal dysbiosis is associated with the loss of the innate anti-HSV activity is likely complex, but our findings suggest that differences in IgG glycan composition contributes as evidenced by the significant reduction in activity when the CVL was treated with PNGase F. We speculate that bacterial glycosidases, which would reduce the affinity for the viral FcγR (gE) (as it does for cellular FcγRs) play a role. This notion is supported by the observation that the three samples from women with CSTIV microbiomes had a predominance of immature glycans detected compared to the three from women with CSTIII (*L. iners* dominant microbiome). Specifically, the latter had higher amounts of bisected N-acetyl glucosamine (GlcNAc) with mono- or di-galactose and no fucose glycans, which have been associated with increased binding affinity for FcγsR (17). However, the three samples from women with *L. crispatus* dominant microbiomes also had a predominance of immature glycans and yet two of these had potent anti-viral activity. These findings suggest a more complex interaction between IgG glycans, the vaginal microbiome and the anti-HSV activity, which will require a larger study.

IgG diffuses rapidly through cervical mucus, accumulates on surfaces and, in a prior study, was shown to trap HSV and prevent infection (25). This trapping was observed at antibody concentrations that were below those needed for neutralization and was lost when the IgG was deglycosylated or when only the Fab was present. Possibly, this trapping was also mediated by interactions between gE and the Fc region of the IgG. However, the IgG Fc-gE interaction is unlikely to be the only mechanism underlying the antiviral activity of genital tract secretions. The antiviral activity was reduced but not abolished when tested against a gE-deleted virus and some antiviral activity was retained in the IgG depleted CVL. This residual antiviral activity could reflect other antiviral proteins such as HNP1-3, which we previously showed has anti-HSV activity, albeit at higher concentrations than detected in the CVL.

In summary, results of these studies provide a potential mechanistic link for the increased risk of HSV infection and replication in the setting of BV or vaginal dysbiosis. Moreover, they demonstrate that the ability of IgG Fc to bind gE may be protective or immune evading, depending on the relative amounts of viral specific antibodies present and whether the IgGs are properly glycosylated. The protective mechanism may also be important for other viruses that express viral FcγRs including human cytomegalovirus (HCMV), which encodes four glycoproteins that act as viral FcγRs and may also play opposing roles in protecting or evading host immunity (26).

## Materials and Methods

### Clinical samples

Stored CVL were available from two prior studies that were conducted in the Bronx, New York and were approved by the Albert Einstein College of Medicine Institutional Review Board; all participants provided written informed consent. CVL was collected by washing with 10 ml of normal saline, divided into aliquots and stored at -80°C. Nugent scores, 16s ribosomal RNA sequencing and concentrations of immune mediators in the CVL were extracted from the data base for each study.(14) HSV serostatus was assessed by HerpeSelect immunoblot assay (FOCUS Technologies, Cypress, CA) in serum.

### Cells and viruses

Vero (monkey kidney) epithelial cell lines were obtained from the American Type Culture Collection. The viruses (HSV-2(G) (27), HSV-1B^3^x1.1 (23) and the HSV-1 gE deleted virus (28) (gift from Richard Roller, University of Iowa) were grown and titered on Vero cells.

### HSV inhibitory activity

CVL was diluted 1:4 in Dulbecco’s Modified Eagle Medium (DMEM) or PBS and mixed with an equal volume of HSV-2(G) virus (∼100-250 pfu) for 1 hour before applying the mixture in duplicate to a confluent monolayer of Vero cells in 24-well plates. After a 1-hour incubation at 37°C, the inoculum was removed by washing and the cells were overlaid with 0.5% methylcellulose in medium 199 supplemented with 1% fetal bovine serum. Controls included HSV-2 mixed with an equal volume of buffer. The number of plaques were quantified after 48 h by crystal violet staining. As indicated, plaque assays were conducted with HSV-1 and gE-deleted virus or with the addition of mouse anti-glycoprotein E (sc-56990, Santa Cruz) or anti-gB (BMPC23) (29) added to the virus prior to infecting cells.

### Enrichment and depletion of immunoglobulins in CVL

Individual or pooled CVL samples were concentrated using Centricon Plus-30 Centifugal filter units (Sigma-Aldrich) and the concentrated material applied to a Protein L column (Thermo Fisher Scientific, Waltham, Massachusetts). The column was washed 5 times and the bound proteins (enriched for immunoglobulins) eluted using 0.1M glycine pH 2.8 buffer and neutralized in 1M Tris, pH 8.8. For depletion studies, the enriched immunoglobulins (eluant from Protein L column) were incubated with either biotinylated goat anti-human IgG (cat # 31770), biotinylated goat anti-human IgA (cat # A18785), or both (Thermo Fisher Scientific, Waltham, Massachusetts). Magnetic streptavidin beads were added for a 2-hour incubation at 4°C and the bound IgA and/or IgG removed with a magnet. The protein concentration in enriched and/or depleted fractions was quantified by nanodrop and material diluted to equivalent protein concentrations before testing for anti-HSV activity in the plaque assay. Commercial enzyme-linked immunosorbent assays (ELISA) were used quantify IgG or IgA in the different fractions (Thermo Fisher Scientific, Waltham, Massachusetts) and, as an additional control, the concentration of human neutrophil peptides 1-3 (HNP1-3) was measure by ELISA (HyCult Biotechnology, Uden, The Netherlands).

### Quantification of HSV-specific antibodies

HSV-specific antibodies in CVL or serum were quantified by ELISA as previously published. (30) Plates were coated with HSV infected and uninfected Vero cell lysates overnight, fixed using 1% paraformaldehyde and permeabilized with 0.1% Triton-X. Plates were then blocked for 2 h and incubated in duplicate with serial dilutions of CVL or serum overnight. Bound antibodies were detected after adding horse radish peroxidase-conjugated anti-human IgG or IgA.

### Glycan analysis

CVL or IgG enriched CVL after depletion of IgA with streptavidin magnetic beads) were deglycosylated by overnight incubation with peptide-N-glycosidase F (PNGaseF) (cat #: NS99010, N-zyme scientific, Doylestown, PA) in PBS at 37°C to release the glycans, which were then precipitated using cold ethanol, and the supernatant containing the released native N-glycans was collected and dried by vacuum centrifugation. The samples were desalted by resuspension in 0.1% trifluoroacetic acid (TFA) and loaded onto graphite spin columns containing porous graphitized carbon (PGC) (cat #: 60106407, Thermo Fisher), washed and eluted from the graphite spin column using 25% acetonitrile/0.1% TFA. Subsequently, the eluted N-glycans were dried by vacuum centrifugation. Permethylation of the native N-glycans was performed by combining the samples with iodomethane (cat #: 289566, Sigma-Aldrich, St. Louis, MO) in anhydrous dimethyl sulfoxide (cat #: 276855, Sigma-Aldrich). The mixture was then loaded onto spin columns containing sodium hydroxide beads (cat #: 367176, Sigma-Aldrich). Purification of the resulting permethylated N-glycans was achieved through liquid-liquid extraction using a 1:1 chloroform/methanol mixture. The chloroform layer, which contained the permethylated N-glycans, was collected and dried under nitrogen gas. The dried permethylated N-glycans were resuspended in 50% methanol. A 1 µl sample was mixed with super-DHB matrix solution (cat #: 63542, Sigma-Aldrich) at a 1:1 ratio and spotted onto an MTP 384 well-polished steel target plate (Bruker Daltonics, Billerica, MA). Analysis of the samples was performed using MALDI-TOF with the Ultraflextreme mass spectrometer (Bruker Daltonics). The acquisition software used was FlexControl 3.4, and the raw data were processed using FlexAnalysis 4.0 software. The resulting mass spectra were converted to peak lists. To determine the percent composition of each N-glycan, the raw abundance of each N-glycan was divided by the sum of the abundances of all N-glycans. All the glycans with m/z above 1836 were considered mature and those below 1836 were defined as immature.

### Statistical Analysis

Unpaired and paired t-tests or ANOVA with multiple comparisons were performed as indicated. Spearman tests were performed to assess correlations between antiviral activity and immune mediators or bacterial concentrations. Analyses were performed using GraphPad Prism, version 10 (GraphPad Software, La Jolla, CA).

## Acknowledgments

This work was supported by R01HD098977, 2R01AI134367, P30 AI124414, UM1 AI068613 and UM1 TR004400. The funders had no role in study design, data collection and interpretation, or the decision to submit the work for publication.

We also thank Richard Roller for the gift of the gE deleted virus and Fereshteh Zandkarimi, Columbia University, for performing mass spectroscopy.

---

**Supplemental Figure S1. HSV-specific IgA quantification in CVL and serum**. HSV-specific IgA was measured via ELISA in serial dilution of CVL (A) and serum (B) in n=9 CVL samples obtained from the control cohort (Table 2). Each participant’s CVL is color-coded.

**Supplemental Figure S2. Intact mAb or Fab of anti-gE, but not anti-gB, contribute to anti-HSV activity in the CVL.** Anti-HSV neutralizing activity was measured in the CVL with increasing concentration of anti-gB (blue) or anti-gE (red) intact mAb (closed symbol, solid line) or Fab fragment (open symbol, dotted line). Results are mean ± sd from duplicate wells.

**Supplemental Figure S3. Individual IgG glycan m/z profile.** IgG glycans of N=9 CVL samples were measured using MALDI-TOF and the relative mass (m/z) and glycan assignment for each sample is shown.

